# Electromagnetic waves destabilize the SARS-CoV-2 Spike protein and reduce SARS-CoV-2 Virus-Like Particle (SC2-VLP) infectivity

**DOI:** 10.1101/2024.09.11.612487

**Authors:** Skyler Granatir, Francisco M. Acosta, Christina Pantoja, Johann Tailor, Angus Fuori, Bill Dower, Henry Marr, Peter W. Ramirez

**Affiliations:** Department of Biological Sciences, California State University Long Beach, Long Beach, CA; Epirus Inc., Torrance, California, USA

**Author notes:** Address correspondence to Peter Ramirez. These authors contributed equally to this work (Skyler Granatir, Francisco M. Acosta, Christina Pantoja).

**Keywords:** SARS-CoV-2, Spike, Electromagnetic Waves, Coplanar Waveguide, Transmission, Sanitation

## Abstract

Infection and transmission of Severe Acute Respiratory Syndrome Coronavirus 2 (SARS-CoV-2) continues to pose a global public health concern. Using electromagnetic waves represents an alternative strategy to inactivate pathogenic viruses such as SARS-CoV-2 and reduce overall transmission. However, whether electromagnetic waves reduce SARS-CoV-2 infectivity is unclear. Here, we adapted a coplanar waveguide (CPW) to identify electromagnetic waves that could neutralize SARS-CoV-2 virus-like particles (SC2-VLPs). Treatment of SC2-VLPs, particularly at frequencies between 2.5-3.5 GHz at an electric field of 400 V/m for 2 minutes, reduced infectivity. Exposure to a frequency of 3.1 GHz decreased the binding of SC2-VLPs to antibodies directed against the Spike S1 subunit receptor binding domain (RBD). These results suggest that electromagnetic waves alter the conformation of Spike, thereby reducing viral attachment to host cell receptors. Overall, this data provides proof-of-concept in using electromagnetic waves for sanitation and prevention efforts to curb the transmission of SARS-CoV-2 and potentially other pathogenic enveloped viruses.

## Introduction

Coronaviruses consist of a family of enveloped positive-sense single-stranded RNA viruses that infect a wide variety of mammals and can cause mild or severe disease^1^. Three human coronaviruses emerged in the 21^st^ century that pose a significant threat to public health: severe acute respiratory syndrome coronavirus 1 (SARS or SARS-CoV-1), middle east respiratory syndrome (MERS) and severe acute respiratory syndrome coronavirus 2 (SARS-CoV-2)^2–5^. SARS-CoV-2 is the causative agent of the COVID-19 pandemic^6^. Viral transmission of human coronaviruses occurs via person-to-person contact, respiratory droplets, and through touching contaminated surfaces^7^. As of August 2024, SARS-CoV-2 has infected nearly 800 million individuals and led to over 7 million deaths (WHO). Despite effective vaccines^8^ and therapeutics^9,10^, SARS-CoV-2 continues to circulate and can lead to complications, particularly in high-risk groups. Therefore, increased strategies to curb SARS-CoV-2 transmission are needed.

Standard pathogen disinfection includes the use of high temperatures, ultraviolet and ionizing radiation, and chemical agents, though these techniques have limitations^11–15^. Electromagnetic waves offer an alternative strategy to inactivate viruses, possessing high penetration, uniform heating, and minimal pollution^16^. How electromagnetic waves alter virus infectivity is an active area of investigation, though it can include changes in virus morphology^15,17^, damage to viral RNA^18^ and denaturation of enveloped virus glycoproteins^19^.

The SARS-CoV-2 Spike glycoprotein mediates attachment and entry into host cells. Spike contains S1 and S2 subunits that exist in a glycosylated trimeric form on the surface of a mature virus. The receptor binding domain (RBD) within the Spike S1 subunit facilitates binding to host cell receptors such as ACE2^5^. A polybasic furin cleavage site is present at the junction between the S1 and S2 subunits^20^. The host protein furin cleaves Spike, exposing an S2’ cleavage site within the S2 domain. Cleavage by the host serine protease TMPRSS2 exposes the S2 fusion peptide (FP), which inserts into the host cell membrane to mediate entry at the cell surface^21,22^. In some cases, SARS-CoV-2 entry can occur via endocytosis whereby Spike cleavage occurs in a pH-dependent manner mediated by the cysteine protease cathepsin L^23^.

The goal of this study was to determine whether electromagnetic waves could affect the infectivity of SARS-CoV-2. For our studies, we adapted a system using SARS-CoV-2 virus-like particles (SC2-VLPs) that package and deliver exogenous RNA transcripts (i.e. luciferase) into target cells^24^. Because SC2-VLPs contain each of the SARS-CoV-2 structural proteins (Spike, Envelope, Matrix and Nucleocapsid), they recapitulate authentic aspects of SARS-CoV-2 entry, assembly and release and are suitable to work with in a Biosafety Level 2 (BSL2) setting. We employed a coplanar waveguide (CPW) to determine the absorption spectrum of SC2-VLPs. We then tested a range of frequencies to determine whether they have any effect on reducing SC2-VLP infectivity. We found that frequencies within 2.5-3.5 GHz reduced infectivity when exposing SC2-VLPs to an electric field of 400 V/m for 2 minutes (180 W input supply to TEM cell). This correlated with reduced binding of antibodies to Spike targeting the S1 RBD, suggesting that electromagnetic waves can induce conformational changes within Spike that negatively impact viral attachment/entry into host cells. To our knowledge, our data is the first to provide direct experimental evidence of using electromagnetic waves to reduce SARS-CoV-2 infectivity and provide a plausible mechanism for how this occurs. This study serves as a proof-of-concept for further development in using electromagnetic waves to combat COVID-19 transmission.

## Results

### Identification of electromagnetic wave frequencies absorbed by SC2-VLPs using a Coplanar Waveguide (CPW)

To explore the permittivity of SC2-VLPs at a range of frequencies, we employed a coplanar waveguide (CPW) in combination with a Vector Network Analyzer (VNA). We modeled our design after Yang et al, who designed a CPW circuit to measure the electromagnetic wave absorption spectra of H3N2 influenza viruses^25^. A CPW is a waveguide in which all conductors supporting wave propagation is placed on the same plane. Active components can be placed along a microstrip (Figure 2A). Each end of the CPW is connected to a VNA, which can emit and receive thousands of frequency points over a short interval of time (Figure 2B). In this manner, the absorption of any fluid along the microstrip can be determined. A higher absorption implies a greater electromagnetic wave permittivity at a given frequency. This can lead to changes in virus architecture, rendering them non-infectious. To obtain an absorption spectrum for SC2-VLPs, we dripped them uniformly along our CPW microstrip (Figure 2A). We observed local peaks around 3.1 GHz and 6 GHz (Figure 2C). It should be noted that even without buffer on the CPW, we had previously observed resonance around 5-6 GHz, likely due to half-wavelength resonance, i.e. half-wavelength would geometrically fit across the CPW at these frequencies and produce a trapped standing wave. Nonetheless, when attempting to remove those contributions, we still witnessed significant absorption by SC2-VLPs at the higher frequencies.

**Figure 1:**
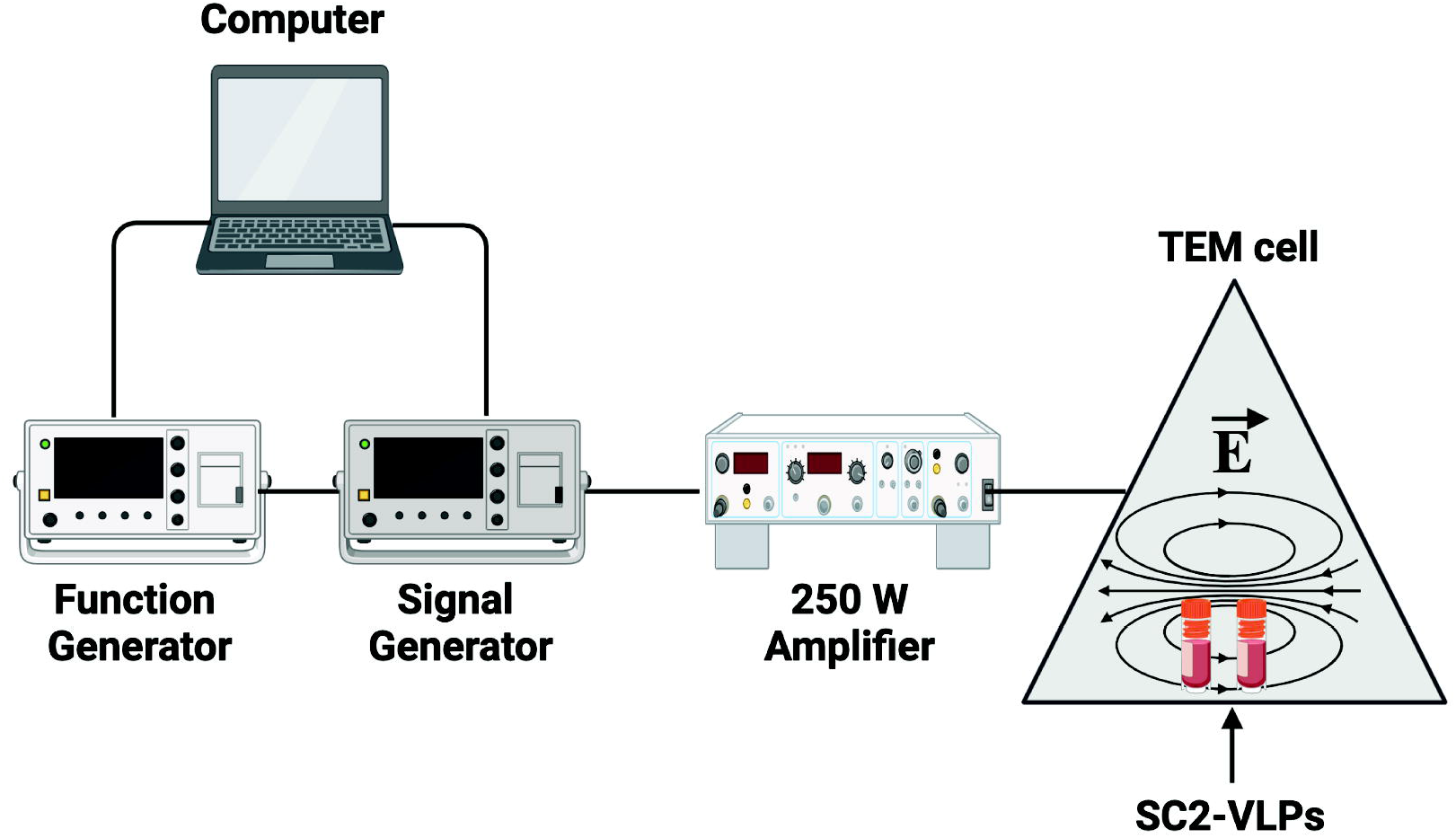
Electromagnetic wave hardware setup. A waveform originates from a signal and function generator commanded by a computer. After the signal generator is triggered by the function generator, the signal is transmitted to a wideband amplifier (AR500M6G). The amplifier outputs 180 W signal to a transverse electromagnetic (TEM) cell through N-type cables.

**Figure 2:**
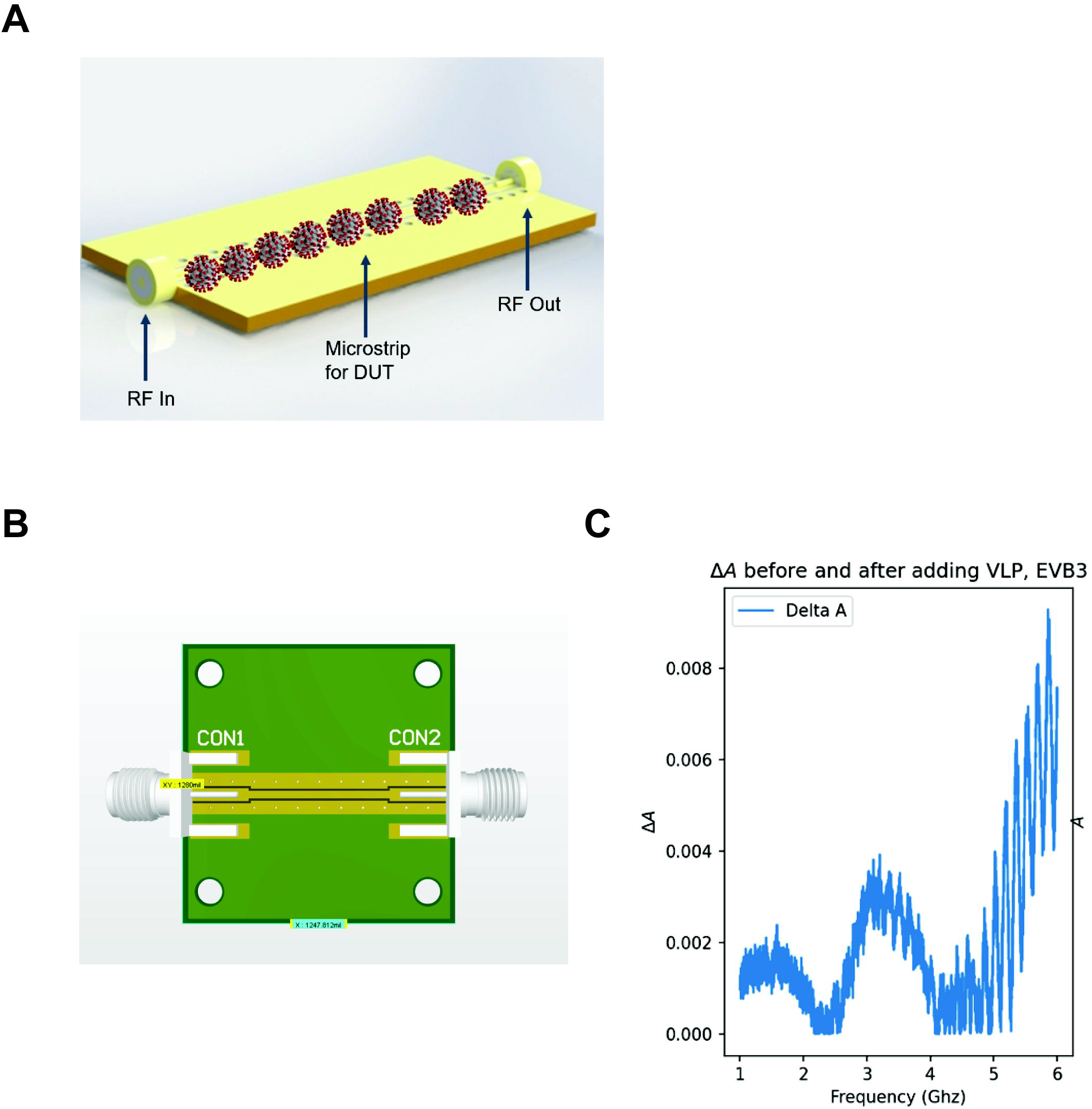
Coplanar Waveguide (CPW) setup and absorption spectra of SARS-CoV-2 Virus-Like Particles (SC2-VLPs). A.) Illustration of a virus suspended solvent on a CPW. B.) Epirus CPW circuit. C.) SC2-VLP absorption spectra. RF: Radio-Frequency signal. DUT: Device-Under-Test

### Electromagnetic waves reduce SARS-CoV-2 Virus Like Particle (SC2-VLP) infectivity

Next, we determined whether exposing SC2-VLPs to a range of electromagnetic frequencies could affect their infectivity (Figure 3A). We chose the following frequencies based on our absorption spectra (Figure 2C): 1.0-2.5 GHz, 2-5-3.5 GHz, 3.5-4.8 GHz, and 4.8-6 GHz. We also treated SC2-VLPs with 2.45 GHz using Microwave (MW) radiation as a control. Overall, treatment of SC2-VLPs with either electromagnetic waves or MW irradiation reduced infectivity relative to untreated samples (Figure 3C). We observed a statistically significant reduction in infectivity when SC2-VLPs were treated with electromagnetic waves within the 2.5-3.5 GHz and 3.5-4.8 GHz range (Figure 3B). We next sought to determine the mechanism for how EMPs reduced SC2-VLP infectivity.

**Figure 3:**
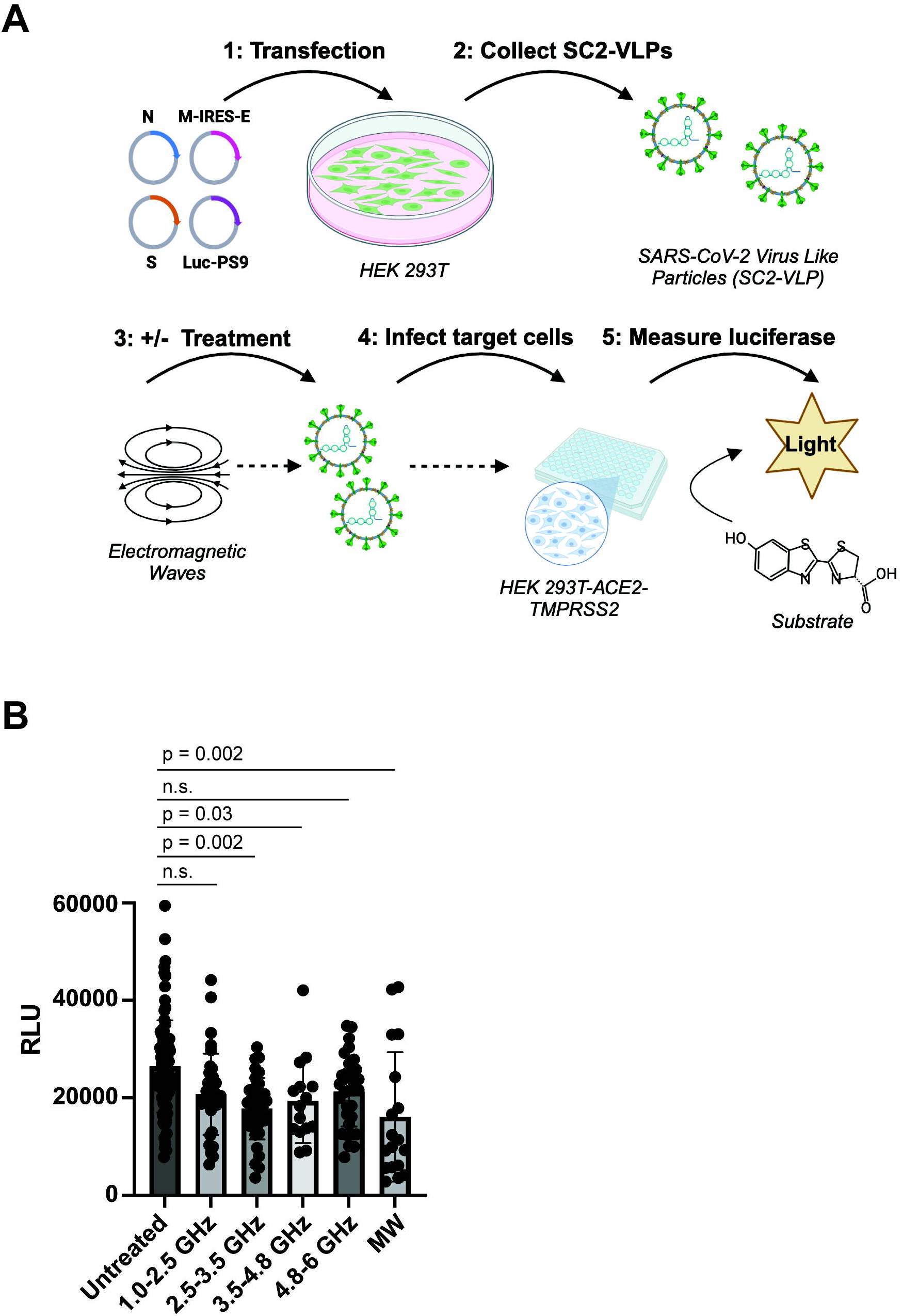
Electromagnetic waves reduce infectivity of SARS-CoV-2 Virus-Like Particles (SC2-VLPs). A.) Diagram of SC2-VLP production, electromagnetic wave treatment and infectivity assay. Plasmids expressing each of the SARS-CoV-2 structural proteins (Nucleocapsid (N); Matrix (M); Envelope (E); and Spike (S)), or a plasmid encoding a packaging signal and luciferase transcript (Luc-PS9) were transfected into viral producer cells (HEK293T). SC2-VLPs were then collected, left untreated or treated with various electromagnetic wave frequencies and used to infect target cells (HEK293T-ACE2-TMPRSS2). The next day, luciferase was measured as a readout of infectivity. Created in BioRender.com. B.) SC2-VLP infectivity assay. Data represent mean +/-SD of 5 independent experiments performed in triplicate. P-values indicate Wilcoxon matched-pair signed rank test of treated compared to untreated samples. (n= 90 (Untreated); n= 36 (1.0-2.5 GHz & 2.5-3.5 GHz) n = 15 (3.5-4.8 GHz), n = 33 (4.8-6 GHz), n = 18 (MW)). RLU: Renilla Luciferase Units (RLU). MW: Microwave

### Electromagnetic waves destabilize the SARS-CoV-2 Spike Receptor Binding Domain (RBD)

Using molecular simulations, applying electric fields induced a conformational change within the recognition loop L3 of the Spike receptor binding domain (RBD), changing two parallel beta sheets into an unstructured coil that diminished binding to ACE2^26^. In another study, applying a 2.45 GHz electromagnetic wave denatured the SARS-CoV-2 Spike S1 subunit *in vitro*^19^. Therefore, we asked whether SC2-VLPs treated with electromagnetic waves led to destabilization of SARS-CoV-2 Spike. We tested this by measuring the ability of untreated or electromagnetic wave treated SC2-VLPs to bind an antibody targeting the Spike S1 receptor binding domain (S1RBD) via Enzyme-Linked Immunosorbent Assay (ELISA) (Figure 4A). We tested electromagnetic wave frequencies that led to either the greatest (3.1 GHz) or smallest (5.9 GHz) reduction in infectivity (Figure 3B). Interestingly, SC2-VLPs treated with 3.1 GHz or 5.9 GHz reduced SARS-CoV-2 Spike binding by approximately 70% and 15%, respectively (Figure 4B). Overall, this data supports a model whereby electric fields alter the conformation of Spike, negatively impacting its ability to bind ACE2 and enter cells.

**Figure 4:**
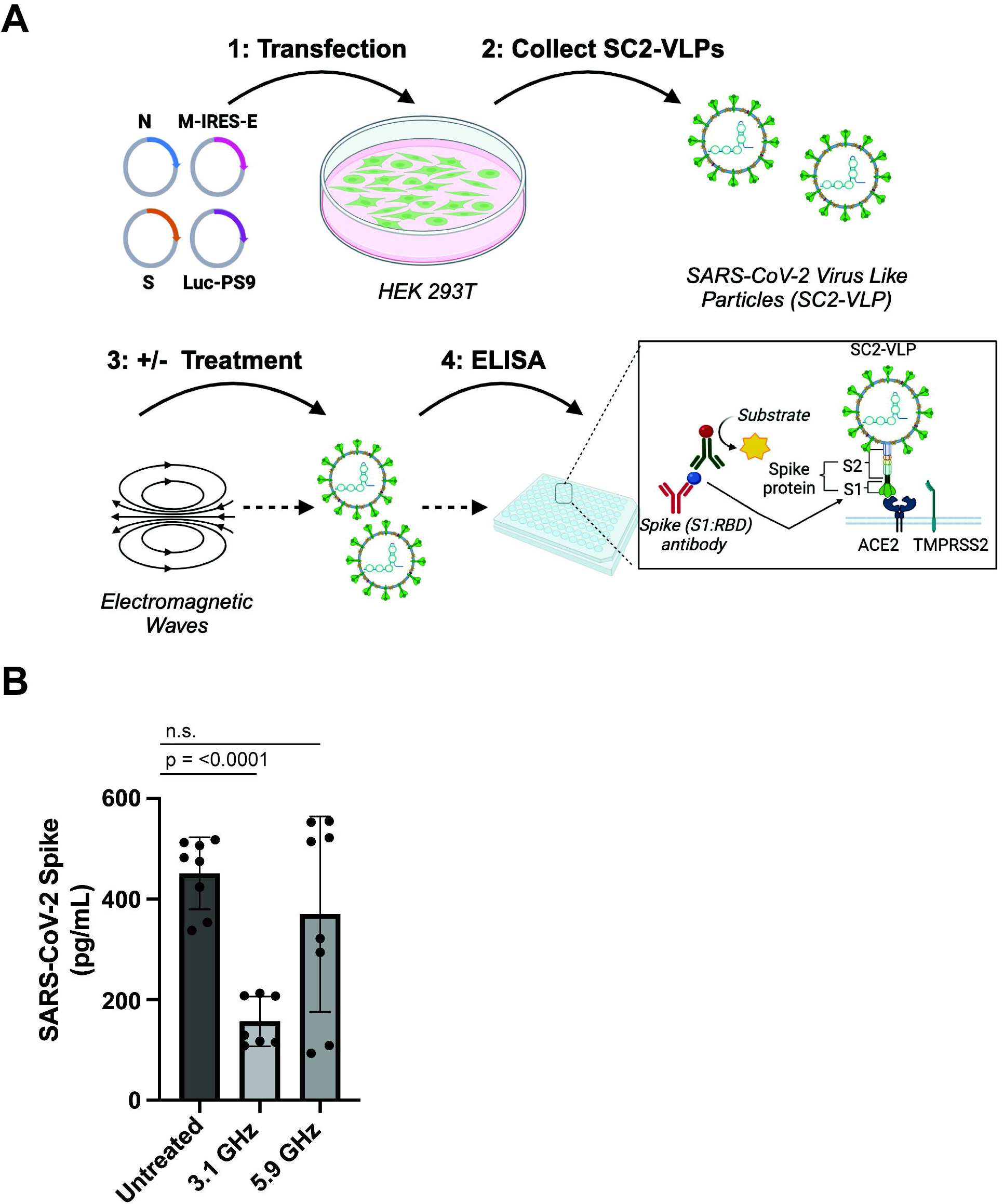
Electromagnetic waves destabilize the SARS-CoV-2 Spike protein. A.) SC2-VLPs were produced, collected, treated with select electromagnetic wave frequencies for 2 minutes, and subjected to an ELISA to measure the stability of the Spike receptor binding domain (RBD). Created in BioRender.com. B.) Spike (S1RBD) ELISA. Data represent +/-SD of 2 independent experiments performed in quadruplicate, except for one condition (3.1 GHz) that was analyzed once in quadruplicate and once in triplicate. P-values indicate one-way ANOVA tests of treated compared to untreated samples.

## Discussion

This study sought to determine the role of using electromagnetic waves to inactivate enveloped viruses, particularly SARS-CoV-2. We present the first proof-of-concept study showing that nonthermal electromagnetic waves of 2.5-3.5 GHz at 400 V/m for 2 minutes alter the conformation of the SARS-CoV-2 Spike S1 RBD, reducing the infectivity of SC2-VLPs.

Mechanistically, inactivation of viruses using electromagnetic waves can occur due to thermal, nonthermal, or physical resonance effects. Thermal effects inactivate viruses by increasing the surrounding temperature. Exposing human coronavirus 229E to electromagnetic waves of 95 GHz at a power density of 70-100 W/cm^2^ for 2 seconds led to virus inactivation induced by a 100° C change in temperature^15^. This resulted in drastic changes to virus morphology, forming holes in the envelope of 229E virions revealed by scanning electron microscopy (SEM)^15^. Siddharta et. al. reduced or completely inactivated Human Immunodeficiency Virus (HIV-1) and Hepatitis C Virus (HCV) using electromagnetic waves at a frequency of 2450 GHz and power densities between 360W – 800W^27^. HIV-1 and HCV inactivation were dependent on the cell culture medium temperature rising from 26° C to 92°C, as no apparent changes in viral infectivity were recorded when the temperature was held constant due to viruses being subjected to either low power (90 or 180 W; 3 minutes) or short-burst high power (600 W or 800 W; 1 minute) electromagnetic waves^27^. A recent study reported a 9.375 GHz frequency at 100 mW/cm^2^ inactivated coronavirus mouse hepatitis A 59 (MHV-A59) by physical destruction of the viral envelope and genome^28^. However, whether these phenomena occurred due to changes in thermal or physical resonance effects were not explored. Here, we did not detect any changes in temperature when subjecting SC2-VLPs to various electromagnetic frequencies (data not shown), suggesting our results occur in a nonthermal manner.

Physical resonance effects occur when objects absorb more energy from their surrounding environment when vibrating at their natural frequency and wavelength. Viruses resonate in the confined-acoustic dipolar mode with electromagnetic waves of the same frequency^16,25^. This leads to a structure resonance energy transfer (SRET) from electromagnetic waves to confined acoustic vibrations (CAV) in viruses, inducing fracture of the virus structure. Indeed, inactivation of influenza A virus strain H3N2 occurred when exposing the virus to a frequency of 6 GHz at 486 W/m^2^ power density, resulting in the physical rupture of the viral envelope through resonance effects despite minimal increases in temperature^25^. We did not directly test whether exposure of SC2-VLPs to 2.5-3.5 GHz led to virus fracture or damaged viral protein(s)/genetic material.

Electromagnetic waves can cause molecules to rotate or vibrate in a nonthermal manner, altering their polarity to induce changes in the conformation of viral proteins. A 2.45 GHz 700W electromagnetic wave denatured the SARS-CoV-2 Spike S1 subunit by 95% in the absence of heat^19^. Thus, conformational changes in Spike occurred due to pure electromagnetic effects. Another study used molecular simulations to conclude that exposure to electric fields can alter the conformation of SARS-CoV-2 Spike and its variants, negatively impacting its ability to bind to ACE2^26^. Interestingly, these simulations reported minimal structural damage to the Spike S2 subunit, suggesting Spike’s susceptibility to electric fields is limited to its prefusion (S1 subunit), rather than post fusion (S2 subunit) conformational state^29^. Our data are suggestive of a nonthermal model, since exposure of SC2-VLPs to distinct electromagnetic waves reduced the binding of Spike to antibodies targeting the S1 RBD (Figure 4B). Future studies should directly test whether the Spike S2 subunit, unlike S1, is relatively resistant to electric fields.

Our study has limitations. First, SC2-VLPs may not wholly reflect what occurs with SARS-CoV-2. Because SARS-CoV-2 is classified as a BSL3 agent, we did not have the research facilities to perform work with this pathogen. However, because SC2-VLPs mimic the viral assembly, packaging, production, and delivery of exogenous transcripts, they provide important insights into the mechanisms governing SARS-CoV-2 replication, including attachment and entry. Nonetheless, future studies should address whether electromagnetic fields can reduce the infectivity of authentic replication-competent SARS-CoV-2. Second, we did not determine whether other SARS-CoV-2 Spike variants are susceptible to electromagnetic fields. Here, we created SC2-VLPs using a plasmid expressing the SARS-CoV-2 Spike protein derived from the original Wuhan-1 strain^24^. Exposure to electric fields via molecular simulations induced structural changes within the SARS-CoV-2 Spike RBDs from wildtype (Wuhan-1) as well as Alpha (B.1.1.7), Beta (B.1.351), Gamma (P.1), Delta (B.1.617.2), and Omicron (BA.1) variants^26,29^. We hypothesize that the structural damage induced by electric fields towards ancestral Spike will extend to other Spike variants. However, this has yet to be experimentally tested. Finally, whether our CPW/VNA system reduces the infectivity of IAV, HIV-1, HCV or other pathogenic enveloped viruses is intriguing but was beyond the scope of this study.

In summary, electromagnetic waves offer an intriguing strategy to inactivate pathogenic viruses. By using a unique CPW and VNA, we rapidly obtained the absorption spectrum of SC2-VLPs, providing insightful information about the permittivity and susceptibility of SARS-CoV-2 to certain electromagnetic wave frequencies. Identifying electromagnetic waves that have minimal impacts on the human body may offer novel *in-vivo* methods for neutralizing pathogenic viruses. Beyond inactivation, the CPW technique employed here could potentially be used to detect viruses with *a priori* knowledge of the virus’s absorption characteristics. Deployment of a portable system utilizing a VNA-like circuit could measure absorption at the most informative frequencies in a wide range of samples. Thus, electromagnetic waves provide several encouraging future developments to improve viral detection, inactivation, and sanitation.

## Materials and Methods

### Hardware

The electronic system consisted of a transverse electromagnetic (TEM) cell, function generator, signal generator, amplifier, and computer (Figure 1). The waveform originated from a signal and function generator commanded by a computer which was interfaced with the equipment to set parameters such as pulse width, input power, and frequency. After the signal generator was triggered by the function generator, the signal was transmitted to a wideband amplifier (AR500M6G), outputting a 180-250 W signal to the TEM cell through N-type cables. The TEM cell used for our experimentation was a Tescom TC-5062C. This machine possessed a shielded enclosure providing powerful electric fields (200 - 2000 V/m). An input power of 180 W, corresponding to 413 V/m, was used for all experiments. Any leaked radiation was below 5 mV/cm^2^, the FCC human safety threshold.

### Cell Lines and Plasmids

HEK293T (ATCC; CRL-3216) were maintained in complete DMEM (Thermo Fisher Scientific): 10% fetal bovine serum (FBS; Corning), glutamine (Thermo Fisher Scientific) and 1% penicillin/streptomycin (Thermo Fisher Scientific). HEK293 cells expressing human ACE2 and TMPRSS2 (Invivogen; hkb-hace2tpsa) were propagated in complete DMEM supplemented with puromycin (0.5mg/ml), hygromycin B (200 mg/ml) and zeocin (100 mg/ml). The following plasmids were gifts from Dr. Jennifer Doudna and purchased from Addgene: CoV-2-N-WT-Hu1 (#177937), CoV-2-M-IRES-E (#177938), CoV-2-Spike-EF1a-D614G-N501Y (#177939), Luc-noPS (#177940), and Luc-PS9 (#177942).

### Generation of SARS-CoV-2 Virus-Like Particles (SC2-VLPs)

HEK293T cells (3.5 × 10^6^ in 10 mL total volume) were seeded in 10 cm dishes 24 hours prior to transfection. The next day, cells were transfected with a total of 20 µg plasmid DNA according to the following molar ratios: *<u>1</u>* CoV2-N-WT-Hu1, *<u>0</u>*.*<u>5</u>* CoV-2-M-IRES-E, *<u>0</u>*.*<u>5</u>* Luc-PS9 or Luc-noPS and *<u>0</u>*.*<u>0125</u>* CoV2-Spike-EF1a-D614G-N501Y. Plasmids were diluted in water to a final volume of 400 µL, along with 100 µL 2.5 M calcium chloride (CaCl_2_) and 500 µL 2X Hepes Buffered Saline (HBS; pH 7.05). This transfection mixture was vortexed, incubated for 1 minute at room temperature and added dropwise over the entire culture dish. To increase transfection efficiency, 10 µL of chloroquine (100 mM) was added prior to returning cells to the incubator (37 °C; 5% CO_2_). Media was changed 16-18 hours post transfection. Forty-eight hours post transfection, supernatant (SC2-VLPs) was collected, clarified by centrifugation, filtered (0.45 µM PES), and stored at -80 °C.

### SC2-VLP infection and luciferase readout

In a clear, sterile, 96-well round-bottom plate, 30,000 HEK293/ACE2/TMPRSS2 cells were either left untreated or mixed with 50 µL SC2-VLPs in triplicate. Cells and SC-VLPs were incubated (37 °C; 5% CO_2_) overnight. The next day, supernatant was removed, cells rinsed with 100 µL 1X PBS (ThermoFisher) and lysed with 20 µL passive lysis buffer (Promega) for 15 minutes at room temperature with gentle rocking. The cells were then transferred to an opaque white 96-well flat-bottom plate, mixed with 50 µL reconstituted luciferase assay buffer (Promega), and luciferase measured immediately on a plate reader (Biotek Synergy H1). The data was expressed as Relative Light Units (RLU).

### Enzyme-Linked Immunosorbent Assay (ELISA)

All reagents, standards, and samples were prepared according to the manufacturer’s instructions (RayBiotech; Catalog #: ELV-COVID19S1). Briefly, 100 uL of diluted standard or sample (untreated or EMP-treated SC2-VLPs) was added to a 96-well plate in duplicate and incubated for 2.5 hours. After incubation, wells were washed and incubated with 100 uL of 1X biotinylated antibody for 1 hour. Following washing, wells were incubated with 100 uL of 1X streptavidin solution for 45 minutes. After a final wash step, 100 uL of a TMB One-Step substrate reagent was added to each well for 30 minutes in the dark, followed by the addition of 50 uL Stop Solution. Absorbance was read immediately at 450 nm on a plate reader (Synergy H1). All incubation steps were carried out at room temperature.

### Data analysis and statistics

Data were analyzed and compiled in Microsoft Excel and GraphPad Prism 9.0 software. Statistical tests were performed in Graph Prism. Significance between samples was assessed using either a Wilcoxon ranked sum test or Analysis of Variance (ANOVA). P values are denoted on the figures. Figures were produced using Adobe Illustrator (CS5) and BioRender software.

## Acknowledgments

We thank Dr. John Guatelli for review and insightful comments on the manuscript. This study was supported, in part, by an NSF SBIR award (#: 2035140) to Epirus, Inc. PWR was supported, in part, by a New Investigator Grant (GF00610932) from the California State University Biotechnology program (CSUBIOTECH) and NIH award R16AI184450. C.P. was supported, in part, by award T32GM138075 from the National Institutes of General Medical Sciences (NIGMS).

## Author Contributions

Conceptualization: S.G., B.D., H.M., P.W.R.; Validation: F.M.A., C.P., S.G., P.W.R.; Methodology, F.M.A., C.P., S.G., J.T., A.F., P.W.R.; Project Administration: P.W.R.; Writing – Original Draft, S.G, F.M.A., P.W.R.; Writing – Review & Editing, S.G, F.M.A., C.P., P.W.R.; Supervision, P.W.R.; Funding Acquisition: H.M., P.W.R.

